# Finely tuned DDA-PASEF-based immunopeptidomics workflow enhances HLA class I coverage

**DOI:** 10.64898/2026.11.03.761330

**Authors:** Jeewan Babu Rijal, Aurélie Hirschler, Arthur Declercq, Margot Sommer, Charline Keller, Arnaud Jung, Alexandre Burel, Luc-Matthieu Fornecker, Delphine Rolland, Ralf Gabriels, Lennart Martens, Christine Carapito

## Abstract

Mass spectrometry-based immunopeptidomics is essential for the direct identification of HLA-presented peptides, but its application to low-input and clinical samples remains challenging. In this study, we present an optimized DDA-PASEF-based workflow to improve HLA class I peptide coverage. First, we optimized a magnetic bead-based sample preparation protocol to support analysis from reduced starting material and increase compatibility with automation. We then evaluated LC gradient length and column dimensions to define complementary configurations for high-throughput and high-coverage analysis. For MS acquisition, we fine-tuned key parameters, including ion accumulation time, number of PASEF scans, ion mobility range, and polygon filtering, to improve sensitivity while maintaining a suitable cycle time and limiting pre-CID truncations. The customized polygon filter improved the detection of singly charged HLA class I peptides, thereby broadening the identified immunopeptide repertoire. Finally, TIMS²Rescore was integrated to improve confidence in peptide identification. Overall, this optimized workflow improves HLA class I immunopeptide coverage across different sample types, even from low injected sample amounts, supporting its future applicability to clinically relevant samples.

## Introduction

The identification of peptides presented by human leukocyte antigen (HLA) molecules, known as immunopeptidomics, is essential for characterizing the antigenic landscape recognized by T cells^1^. This promising approach plays a central role in the discovery of clinically relevant antigens for cancer immunotherapy, infectious disease monitoring,^2^ and personalized vaccine development^3^. Mass spectrometry (MS) is uniquely suited for immunopeptidomics because it enables the unbiased identification of naturally presented peptides isolated from HLA^4,5^, including post-translationally modified peptides and mutation-derived neoepitopes.^6^

Despite its potential, immunopeptidomics remains technically challenging across the entire workflow. HLA complexes are immunoaffinity-purified from cell or tissue lysates, followed by acidic elution of their bound peptides. Consequently, the workflow is susceptible to residual contaminants, such as polyethylene glycol (PEG)^7^ and detergents, which can impair chromatographic separation and ionization efficiency. High analytical sensitivity is a prerequisite to achieve sufficient coverage for the detection of rare but clinically important antigens^8,9^. Moreover, the absence of defined proteolytic cleavage rules^10^, the large search space of potential peptide sequences, and the low signal intensity of many HLA ligands make immunopeptidomics data interpretation challenging and increase the risk of false-positive and false-negative identifications^11,12^.

Historically, conventional protocols have required tens to hundreds of millions of cells or large tissue sections to generate sufficient material for analysis, limiting their broader applicability to clinically relevant samples such as tumor biopsies and scarce immune cell populations^13^. Recent advances have addressed individual components of this workflow. Magnetic bead-based and semi-automated enrichment protocols have improved scalability, while TIMS-based PASEF acquisition has increased sensitivity and enabled tailored precursor selection in the ion mobility–m/z space. For HLA class I peptides, the inclusion of singly charged precursors and prediction-assisted rescoring can further increase identification coverage.^14–16^ However, these developments have largely focused on individual components of the workflow, and an integrated evaluation combining sample preparation, chromatographic separation, DDA-PASEF acquisition parameters, and premature peptide fragmentation occurring before the intended collision-induced dissociation step, hereafter referred to as pre-CID truncation, remains lacking for the timsTOF Ultra platform.

Here, we systematically evaluated three major components of the workflow: (i) the adaptation and optimization of an HLA class I sample preparation protocol using MagReSyn® Protein A magnetic beads for reduced starting material and increased compatibility with automation, in combination with the addition of the MS-compatible nonionic surfactant DDM; (ii) the evaluation of nanoLC separation and optimization of DDA-PASEF acquisition on the timsTOF Ultra 2; and (iii) the integration of TIMS²Rescore^17^ to incorporate predicted peptide properties into identification confidence assessment. Rather than defining a single universally optimal method, we aimed to identify configurations suited to different requirements for sensitivity, throughput, and immunopeptidome coverage.

## Results

### Optimisation of a magnetic bead-based sample preparation protocol for reduced starting material

We first compared two sample preparation protocols using HL60 cells. A commonly-used sepharose-based protocol was performed using 600 million starting cells, whereas 60 million cells were used for the magnetic bead-based protocol. HLA peptides corresponding to 30 million and 15 million cell equivalents were subsequently injected, respectively. Despite the 10-fold lower starting material and two-fold lower injected cell equivalent, the magnetic bead protocol identified 20,109 peptides compared with 17,682 peptides using sepharose beads, corresponding to an approximately 14% increase in peptide identifications (**Fig. 2A**).

**Fig. 1.**
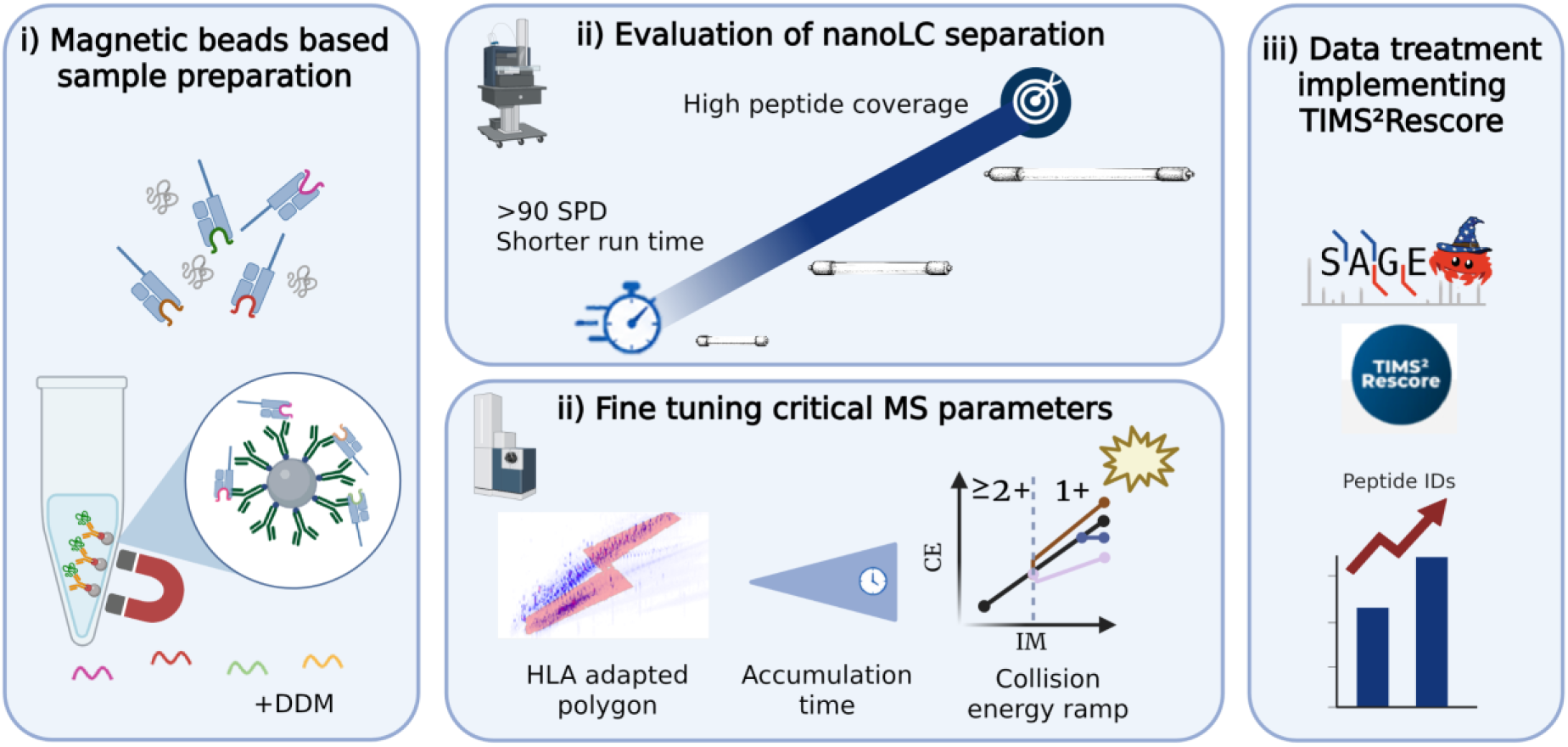
Immunopeptidomics workflow optimisations. (i) Adaptation and optimization of an HLA class I sample preparation protocol using MagReSyn® Protein A magnetic beads for reduced starting material and increased compatibility with automation; (ii) evaluation of nanoLC separation and optimization of DDA-PASEF acquisition on the timsTOF Ultra 2; and (iii) integration of TIMS²Rescore to incorporate predicted peptide properties into identification confidence assessment.

**Fig. 2.**
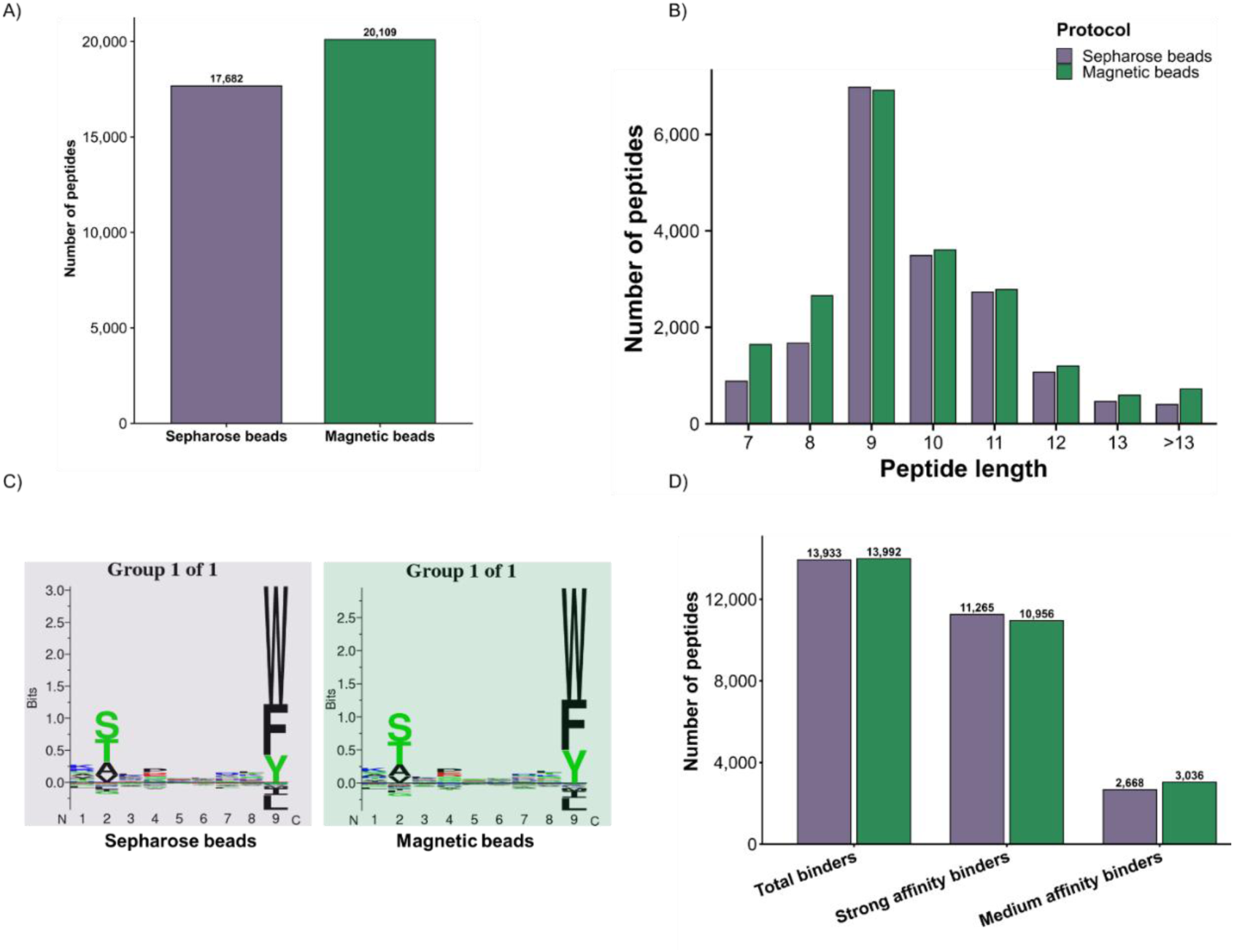
Comparison of sepharose bead-and magnetic bead-based immunopeptidomics protocols. HLA peptides corresponding to 30 and 15 million HL60 cell equivalents were injected for the sepharose bead- and magnetic bead-based protocols, respectively, using a 60-min LC gradient. A) Total number of identified peptides. B) Peptide length distribution C) GibbsCluster 2.0 sequence motifs generated from identified 8-13-mer peptides D) Number of total, strong-affinity, and medium-affinity HLA binders among 8-13-mer peptides predicted using NetMHCpan 4.1.

The identified peptides showed comparable length distributions between the two protocols, with 9-mers representing the predominant population (**Fig. 2B**). GibbsCluster^18^ analysis also revealed similar HLA-I sequence motifs (**Fig. 2C**). NetMHCpan 4.1^19^ predictions identified 13,933 and 13,992 total HLA binders for the sepharose and magnetic bead protocols, respectively, with comparable numbers of strong-and medium-affinity binders (**Fig. 2D**). Based on these results and its performance with reduced starting material, the magnetic bead-based workflow was selected for subsequent optimization at lower sample inputs.

### Optimisation of LC chromatographic separation

#### A 5 cm column enables high-throughput immunopeptidome analysis

We evaluated seven different LC gradient lengths (10 to 150 min) and three analytical columns (5, 25, 50 cm) to determine conditions suitable for high-throughput and high-coverage immunopeptidomics. Detailed LC gradients used are provided in **Supplementary Table 1**. We first compared 5 and 25 cm Aurora (IonOpticks) columns using short LC gradients. With a 10-min gradient, the 5 cm column identified 5,146 peptides compared with 4,098 peptides using the 25 cm column (**Fig. 3A**). Increasing the gradient to 15 min resulted in 7,863 and 7,041 peptide identifications, respectively. At 20 min, both columns provided nearly identical coverage, with 9,649 peptides identified using the 5 cm column and 9,588 using the 25 cm column. With a 30-min gradient, however, the 25 cm column provided higher coverage than the 5 cm column, identifying 12,942 compared with 10,360 peptides. Thus, the 5 cm column was particularly advantageous for very short gradients, allowing more than 5,100 peptides to be identified with only a 10-min active gradient and enabling a practical throughput of up to 90 samples per day.

**Fig. 3.**
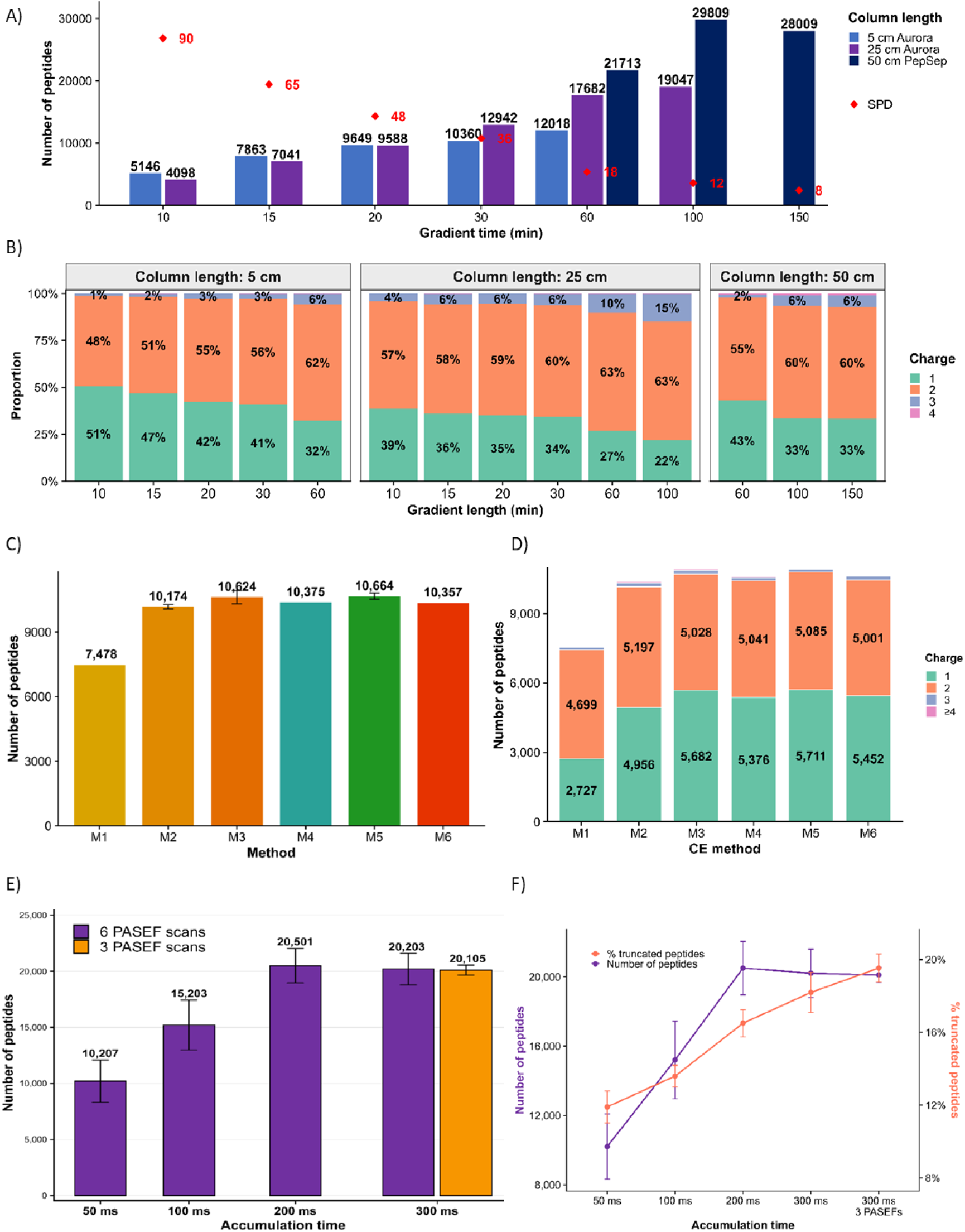
Fine tuning critical LC and MS parameters. A) Number of identified peptides using different gradient lengths (10-150 min) and column lengths (5, 25 and 50 cm in blue, light purple, dark purple, respectively) following injection of HLA peptides corresponding to 30 million HL60 cell equivalents. The red diamonds show the calculated samples per day for each gradient lengths B) Proportion of peptide ions’ charge states observed for the different column lengths and their respective LC gradients. C) Number of peptides identified using different collision energy ramps following injection of HLA peptides corresponding to 30 million HL60 cell equivalents and a 20-min LC gradient. D) Number of identified peptide ions according to charge state for the different collision energy ramps. E) Number of peptides identified using accumulation times ranging from 50 to 300 ms and either 6 PASEF scans (purple) or 3 PASEF scans (orange), following injection of HLA peptides corresponding to 5 million HL60 cell equivalents and a 100-min LC gradient. F) Number of identified peptides and proportion of pre-CID truncated peptides across the different accumulation time and PASEF scan conditions.

#### A 50 cm column provides the highest immunopeptidome coverage

To maximize peptide coverage, we subsequently evaluated longer gradients using 25 cm Aurora (IonOpticks) and 50 cm PepSep (Bruker) columns. With the 50 cm column, peptide identifications increased from 21,713 using a 60-min gradient to 29,809 using a 100-min gradient, representing the highest coverage obtained in this experiment (**Fig. 3A**). Extending the gradient further to 150 min did not provide an additional benefit, with 28,009 peptides identified. Therefore, the combination of a 50 cm PepSep column and a 100-min gradient enabled maximum immunopeptidome depth.

#### Shorter LC gradients result in a higher proportion of singly charged peptides

The precursor charge-state distribution also varied according to gradient length (**Fig. 3B**). Across the different column lengths, shorter gradients consistently resulted in a higher proportion of singly charged peptides, whereas the relative contribution of doubly and triply charged peptides increased with longer gradients. For the 5 cm column, singly charged peptides represented 51% of identifications with the 10-min gradient and progressively decreased to 32% with the 60-min gradient, while doubly charged peptides increased from 48% to 62%. A similar trend was observed with the 25 cm column, where singly charged peptides decreased from 39% with a 10-min gradient to 22% with a 100-min gradient, accompanied by an increase in doubly and triply charged peptides. With the 50 cm column, singly charged peptides decreased from 43% at 60 min to 33% at both 100 and 150 min.

Overall, these experiments defined two complementary LC conditions depending on the analytical objective. A 5 cm Aurora column with a 10-min gradient provided the highest throughput (90 SPD) while maintaining substantial peptide coverage, whereas a 50 cm PepSep column with a 100-min gradient provided the highest overall immunopeptidome depth at a throughput of 12 SPD.

### Fine-tuning critical MS parameters on timsTOF Ultra

We initially employed a method previously optimized on the timsTOF Pro 2 (**Suppl. Fig. 1**) as the starting point for systematically optimizing critical MS acquisition parameters specifically for the timsTOF Ultra 2.

#### Collision-energy optimization preferentially improves singly charged peptide identification

We evaluated six different linear and non-linear CE profiles (M1 to M6), as illustrated in **Suppl. Fig. 2**, using the ion mobility range previously optimized on the timsTOF Pro 2 (0.6–1.7 1/K₀). Particular attention was given to the higher ion mobility region (1.2–1.7 1/K₀), where a large proportion of singly charged HLA-I peptide precursors are detected (**Fig. 3C–D**).

Total peptide identifications were similar across methods M2–M6, ranging from 10,174 to 10,664 peptides, whereas the lower-energy M1 method resulted in only 7,478 identifications (**Fig. 3C**). Inspection of the precursor charge-state distribution showed that this difference was predominantly associated with singly charged peptides. M1, which applied the lowest CE in the 1.2–1.7 1/K₀ region, identified 2,727 singly charged peptides, compared with 5,711 using M5, corresponding to more than a 2-fold increase. In contrast, doubly charged peptide identifications increased only from 4,699 to 5,085 (+8%) between M1 and M5 (**Fig. 3D**). Thus, insufficient collision energy particularly impaired the identification of singly charged HLA peptides, whereas doubly charged precursors were comparatively less affected.

Among the higher-energy profiles, M5, in which CE increased linearly from 20 to 59 eV between 0.6 and 1.6 1/K₀ and was maintained at 59 eV between 1.6 and 1.7 1/K₀, provided the highest mean peptide coverage. However, the improvement in total identifications was marginal compared with M3 (10,664 versus 10,624 peptides; <1%). Importantly, however, M5 also showed improved reproducibility, with the relative standard deviation decreasing from approximately 3% for M3 to 1% for M5. We therefore selected the M5 CE profile for subsequent method optimization, based on its combination of high peptide coverage, efficient recovery of singly charged precursors, and improved reproducibility.

#### Increased accumulation time improves the immunopeptidome coverage

Building on our previous optimization on the timsTOF Pro 2 **(Suppl Fig. 1)**, we further evaluated the effect of TIMS accumulation time on immunopeptidome coverage using the timsTOF Ultra 2. Accumulation times ranging from 50 to 300 ms were tested with 6 PASEF scans. At 300 ms, we additionally reduced the number of PASEF scans from 6 to 3 to determine whether shortening the acquisition cycle time could compensate for the longer accumulation time.

Increasing the accumulation time from 50 to 200 ms markedly improved peptide coverage, with the number of identified peptides increasing from 10,207 to 20,501, corresponding to an approximately two-fold increase (**Fig. 3E**). Increasing the accumulation time further to 300 ms did not provide additional benefit, with 20,203 peptides identified, indicating that the gain in sensitivity had reached a plateau under these conditions. Longer TIMS accumulation times also increased the total acquisition cycle time **(Suppl. Table 2)**, with the 300-ms method exceeding 2 s. We therefore reduced the number of PASEF scans from 6 to 3 at 300 ms, bringing the cycle time below 2 s. This modification slightly improved the repeatability of peptide identifications, as reflected by the reduced variability between replicates, but did not increase overall peptide coverage, with 20,105 peptides identified compared with 20,203 using 6 PASEF scans (**Fig. 3E**). These results indicate that, on the timsTOF Ultra and under the chromatographic conditions used here, the sensitivity benefit of increasing accumulation time had already plateaued at approximately 200 ms, and shortening the cycle time at 300 ms of accumulation using 3 PASEF scans was insufficient to provide additional identifications.

The proportion of pre-CID truncated peptides also progressively increased with accumulation time (**Fig. 3F**). Between 50 and 200 ms, peptide identifications increased approximately two-fold, whereas the proportion of pre-CID truncated peptides increased by only approximately 4%. Thus, the gain in peptide coverage clearly outweighed the accompanying increase in precursor fragmentation up to 200 ms. Beyond 200 ms, however, the proportion of pre-CID truncated peptides continued to increase without a corresponding improvement in peptide identification. Together, these observations strongly support selecting a 200 ms accumulation time with 6 PASEF scans, which provides the highest peptide coverage while maintaining a shorter acquisition cycle and limiting further increases in pre-CID fragmentation.

Based on the combined optimization, M5, the method using a CE ramp increasing linearly from 20 to 59 eV over the 0.6–1.6 1/K₀ range, followed by a constant CE of 59 eV up to 1.7 1/K₀, together with an accumulation time of 200 ms and 6 PASEF scans, was selected for subsequent method development.

#### TIMS²Rescore increases HLA-I immunopeptide identifications while preserving repertoire characteristics

To evaluate whether data-driven rescoring could further increase immunopeptide identification, Sage search results were rescored using TIMS²Rescore across the complete HL60 dilution series. TIMS²Rescore consistently increased the number of identified peptides across all injected cell equivalents (**Suppl. Fig. 4A**), with a substantial fraction recovered exclusively after rescoring (**Suppl. Fig. 4B**). Importantly, these additional peptides retained the characteristic HLA-I length distribution, dominated by 9-mers (**Suppl. Fig. 4C**), and showed a comparable GibbsCluster motif to Sage-only peptides (**Suppl. Fig. 4D**). Thus, TIMS²Rescore increased immunopeptide coverage while preserving the expected characteristics of the HLA-I repertoire.

#### High immunopeptidome coverage is retained at low injected cell equivalents

We next evaluated the sensitivity of the optimized workflow using a serial dilution of an HL60 immunopeptide sample, corresponding to 15 million down to 0.25 million cell equivalents injected. Peptide identifications remained high even at substantially reduced sample amounts (**Fig. 4A**). Indeed, 5,807 peptides were identified from 1 million cell equivalents injected, while 1,467 peptides were still identified from only 0.25 million cell equivalents. Increasing the injected amount to 5 million cell equivalents markedly increased the number of identifications to 12,297 peptides. Peptide coverage increased further to 15,960 and 15,857 peptides at 10 and 15 million cell equivalents, respectively, indicating that identifications approached a plateau at the highest injected amounts. The proportion of pre-CID-truncated peptides progressively decreased with sample input, from 30% at 15 million cell equivalents to 8% at 0.25 million cell equivalents.

**Fig. 4.**
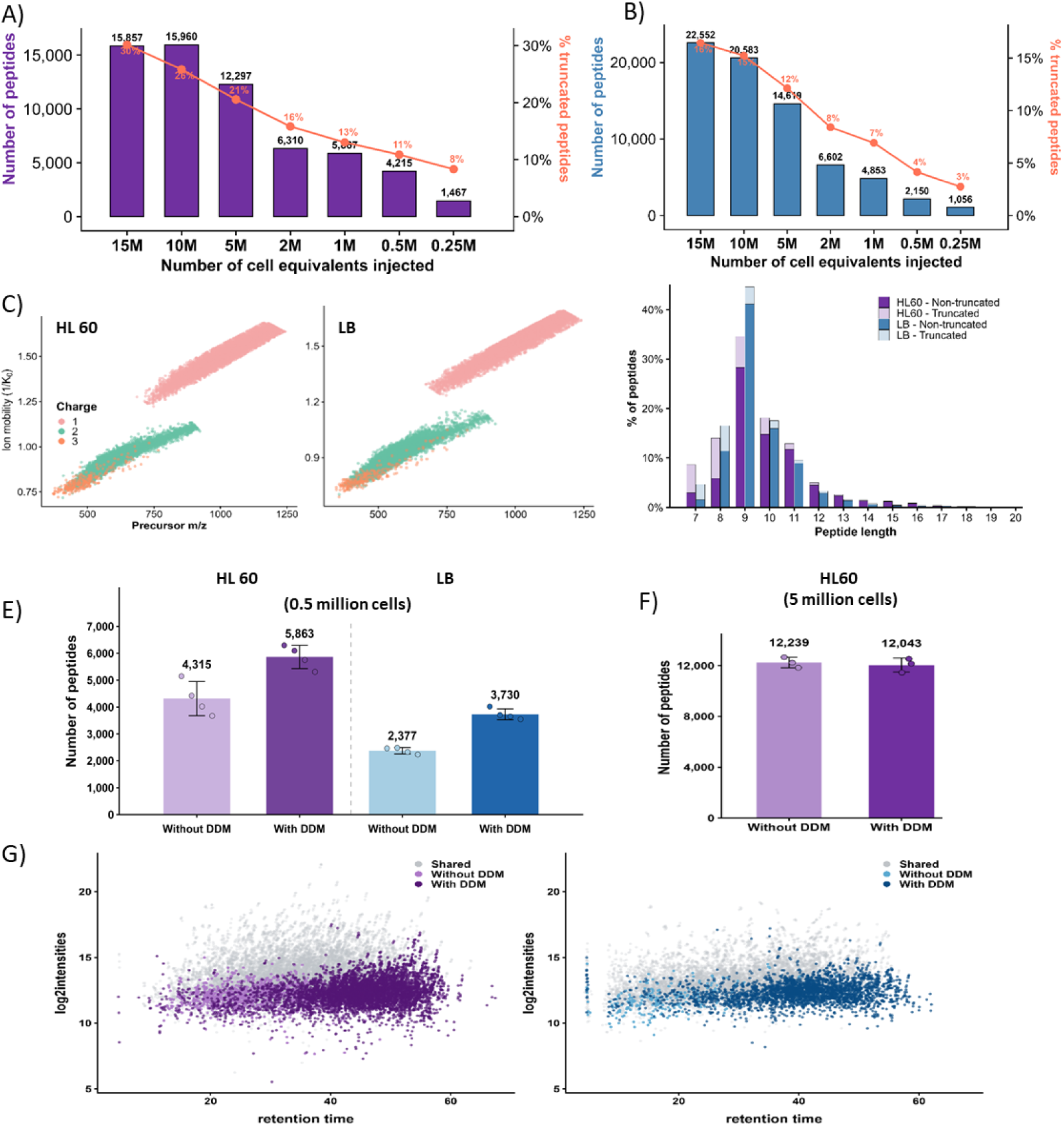
High immunopeptide coverage across decreasing sample inputs. Number of identified peptides and proportion of pre-CID truncated peptides following injection of HLA peptides corresponding to 0.25-15 million cell equivalents using a 25 cm column and a 60-min LC gradient from A) HL60 cells and B) B lymphocytes. The proportion of pre-CID truncated peptides is shown in orange. C) Distribution of identified peptide precursors in the ion mobility (IM) versus m/z space for HL60 and B lymphocytes. D) Peptide length distribution of identified peptides from HL60 cells and B lymphocytes, together with the proportion of pre-CID truncated peptides. For panels C and D, HLA peptides corresponding to 10 million cell equivalents were used for both HL60 cells and B lymphocytes. E) Number of identified peptides without (light color) or with 0.015% DDM (dark color). HLA peptides corresponding to 0.5 million cell equivalents were used for both HL60 cells (purple) and B lymphocytes (blue). F) Number of identified peptides from 5 million HL60 cells equivalents, without (light purple) or with 0.015% DDM (dark purple). G) MS1 peptides intensity profiles in HL60 (left) and LB cells (right) with and without DDM for 0.5 million cells equivalent injections. Log2-transformed MS1 intensities are plotted against retention time (min), pooling data from four replicate injections per condition. Shared peptides are shown in grey; peptides identified exclusively without or with DDM are shown in light or dark colours, respectively.

To assess the transferability of the workflow to another sample type, an HLA class I-enriched LLC B-lymphocyte sample was analysed over the same input range using a timsTOF Ultra equipped with the latest AIP configuration (**Fig. 4B**). Between 1,056 and 22,552 peptides were identified from 0.25 and 15 million cell equivalents, respectively. The proportion of pre-CID truncated peptides similarly decreased from 15% to 2% across this dilution series. The peptide distributions in the ion mobility-m/z plane were comparable between HL60 cells and B lymphocytes, supporting the transferability of the polygon filter to the B-lymphocyte sample (**Fig. 4C**). Both sample types also exhibited similar peptide-length profiles, dominated by 8–11-mer peptides, with 9-mers being the most abundant (**Fig. 4D**). Nevertheless, further optimization specifically for the AIP configuration may provide additional improvements, particularly at the lowest sample inputs. Overall, these results demonstrate that the optimized workflow provides substantial immunopeptidome coverage across different sample types, even from low injected sample amounts.

To further improve peptide recovery at low sample amounts, we evaluated the addition of the MS-compatible nonionic surfactant n-dodecyl-β-D-maltoside (DDM). At 0.5 million injected cell equivalents, DDM increased the number of identified peptides from 4,315 to 5,863 in HL60 samples, corresponding to a 35% increase, and from 2,377 to 3,730 in LB samples, corresponding to a 57% increase (**Fig. 4E**). In contrast, when evaluated at a higher input of 5 million HL60 cell equivalents, DDM provided no additional benefit, with 12,239 and 12,043 peptides identified without and with DDM, respectively (**Fig. 4F**). The MS1 intensity profiles for 0.5 million injected HL60 or LB cells show that peptides gained with the addition of DDM are mostly hydrophobic and of low intensities (**Fig. 4G**). These results indicate that, under our experimental conditions, the benefit of DDM was predominantly observed at low peptide input and for low abundant hydrophobic peptides.

## Discussion

In this study, we systematically optimized a DDA-PASEF-based complete workflow for HLA class I immunopeptidomics, including immunoaffinity purification, chromatographic separation, precursor cloud selection, collision energy, TIMS accumulation time, number of PASEF scans and computational rescoring. Rather than defining a single universally optimal method, our results establish complementary configurations that can be selected according to the available sample amount, required analytical throughput, and desired immunopeptidome depth. This distinction is important for clinical immunopeptidomics, where sample availability and analysis time may be as limiting as instrumental sensitivity.^3^

Conventional immunopeptidomics workflows using CNBr-activated Sepharose beads often require substantial amounts of starting material and involve numerous manual steps, limiting their applicability to low-input and automation. To overcome these limitations, we adapted and optimized a magnetic bead-based SAPrIm protocol¹⁴, substantially reducing material requirements while enabling integration with robotic platforms.

Despite using 1/10th the number of cells for sample preparation and injecting half the cell-equivalent amount, the magnetic bead based protocol yielded similar peptide identifications as the conventional workflow. The expected predominance of 9-mer peptides, the HLA-binding motifs, and the numbers of predicted strong and medium affinity binders were maintained. Therefore, the improved peptide recovery did not appear to introduce a noticeable bias in the canonical HLA class I peptide repertoire. Importantly, the similar number of predicted HLA binders obtained with each of the two protocols indicates that the principal benefit of the magnetic bead workflow is not simply an increase in all identification categories, but the preservation of substantial canonical HLA-I coverage from considerably less starting and injected material.

These findings are consistent with the rationale underlying SAPrIm, which introduced magnetic Protein A beads and semi-automated processing to increase sample throughput and reduce dependence on large amounts of biological material^14^. Magnetic supports facilitate reproducible bead handling, washing, and automation, while reducing the number of manual transfer steps at which low-abundance peptides may be lost. Recent developments have further emphasized the importance of integrating low-volume immunoaffinity capture, automated washing, and peptide purification to combine sensitivity with throughput^20^. The present results extend this direction by showing that improved sample preparation remains beneficial when combined with acquisition specifically tailored for the physicochemical properties of HLA class I peptides.

Nevertheless, the comparison should be considered an end-to-end workflow assessment rather than direct comparison of protocols, as starting cell numbers, injected cell equivalents, and processing steps differed. Thus, the observed improvement cannot be attributed only to the bead material. This is important because immunoprecipitation procedures can influence both peptide yield and repertoire composition, including peptide hydrophobicity^5^. The preservation of peptide-length distributions, binding motifs, and predicted binders in our study argues against a major bias. However, further comparisons with matched inputs, more replicates, and diverse biological matrices are needed to determine which workflow modifications drive the improvement.

An additional consideration for low-input workflows is peptide loss through nonspecific adsorption to sample tubes and other surfaces, which becomes increasingly important as peptide amounts decrease. The MS-compatible nonionic surfactant n-dodecyl-β-D-maltoside (DDM) has been shown to reduce surface-associated peptide losses and improve peptide recovery in low-input proteomics, particularly for hydrophobic peptides^21,22^. More recently, addition of 0.02% DDM to the peptide reconstitution solution substantially increased peptide and protein identifications at single-cell-equivalent sample loads on the timsTOF SCP^23^. Although the potential utility of DDM has previously been proposed for immunopeptidomics, its benefit in this context has not yet been established. Indeed, a recent low-input immunopeptidomics study observed no improvement following DDM addition^24^, whereas an independent conference study reported improved HLA-I peptide recovery when DDM was included during peptide elution, particularly at low PBMC inputs^25^. Our results provide the first direct demonstration of a beneficial effect of DDM on immunopeptide recovery, especially for low abundant hydrophobic peptides. Those results provide evidence that reduction of surface-associated peptide losses can be beneficial in low-input immunopeptidomics. These findings suggest that the effect of DDM may depend on where it is introduced in the workflow and on the amount and nature of the peptide material being analyzed, and support its consideration as a simple strategy for improving sensitivity when sample availability is limiting.

Moving forward, chromatographic optimization revealed two distinct operating regimes, highlighting a fundamental trade-off between depth and throughput and demonstrating that neither can be optimized independently.

The observed charge-state distributions provide additional information beyond total peptide numbers. Across all column lengths, shorter gradients produced a higher proportion of singly charged identifications, whereas the relative contributions of doubly and triply charged precursors increased with longer gradients. This likely reflects changes in chromatographic co-elution, ionization conditions at the time of elution and precursor competition for DDA selection. With steeper gradients, multiply charged precursors may experience greater competition within more densely populated region of the ion mobility-m/z space. Longer gradients reduce co-elution and may therefore allow a larger proportion of these precursors to be selected and successfully identified. Singly charged HLA-I peptides occupy a more distinct region at the ion mobility dimension and may remain comparatively accessible under highly compressed chromatographic conditions.

The importance of singly charged precursors is one of the major difference between immunopeptidomics and conventional tryptic proteomics. HLA-I peptides are generally short and non-tryptic, and a substantial subset may be detectable only as singly charged ions. Standard proteomics acquisition polygons that primarily target multiply charged tryptic peptides can therefore systematically exclude part of the immunopeptidome. Thunder-DDA-PASEF addressed this limitation by using an HLA-I-tailored isolation polygon covering the relevant singly and multiply charged precursor populations.^16^ Our results here reinforce this strategy and further show that inclusion within the acquisition polygon is not sufficient by itself, but the collision energy applied to the corresponding ion mobility region must also produce informative MS/MS spectra.

Collision-energy optimization had a particularly strong effect on singly charged HLA-I peptides and represented the population that was the most sensitive to insufficient collision energy. Because collision energy in timsTOF methods is commonly defined as a function of ion mobility, this result also demonstrates that a conventional mobility-dependent ramp optimized around multiply charged tryptic peptides cannot be assumed to be suitable for singly charged immunopeptides.

The greater sensitivity of singly charged peptides to collision energy is mechanistically plausible, as peptide-fragmentation efficiency depends on precursor m/z, charge state, sequence, and proton mobility^26,27^. Measurements of large collections of synthetic non-tryptic peptides have further demonstrated that collision energy strongly affects the information content of timsTOF MS/MS spectra and that instrument-specific fragmentation models improve spectral prediction^28^. The integration of timsTOF-specific fragmentation and ion-mobility predictions into data-driven rescoring has also been shown to improve the interpretation and identification of peptides from DDA-PASEF spectra^17^. Our findings provide an experimental counterpart to these computational observations that inadequate collision energy does not merely change fragment-ion intensities but can prevent successful identification of a substantial precursor subpopulation.

TIMS accumulation time represented a second major determinant of sensitivity. Increasing the accumulation time from 50 to 200 ms approximately doubled peptide coverage, demonstrating that a longer accumulation period substantially improves the probability of detecting and fragmenting low-abundance immunopeptides. This benefit is consistent with the dual-TIMS design, in which ions are accumulated while a previously accumulated ion population is mobility-separated and analyzed. PASEF thereby combines a high ion utilization efficiency with rapid precursor sequencing^29^. This architecture is particularly advantageous for immunopeptidomics because the isolated HLA peptide mixture contains many low-abundance precursors distributed across a wide range of sequences and physicochemical properties.

The optimal accumulation time identified here differs from the 300-ms ramp combined with three MS/MS frames reported for Thunder-DDA-PASEF on the timsTOF Pro 2^16^. This difference should not be interpreted as a contradiction. The two studies used different instrument generations, samples, chromatographic conditions, and acquisition configurations. The timsTOF Ultra also incorporates a more compact ion-transfer and TIMS configuration than the timsTOF Pro platform. This architectural difference may alter the relationship among ion residence time, transmission, capacity, and precursor fragmentation. However, the effect of cartridge geometry was not evaluated experimentally in the present study and should therefore be considered a possible rationale. More broadly, the difference between the optimal settings emphasizes that parameters established on one timsTOF model should be re-evaluated before transfer to another platform.

Pre-CID fragmentation can generate N- or C-terminally truncated products from a genuine HLA ligand before its intended MS/MS fragmentation. These products may subsequently be selected as independent precursors and assigned as distinct peptide sequences, thereby inflating the apparent diversity through sequences differing by one or more terminal residues. This issue is particularly relevant in immunopeptidomics because database searches are performed without enzyme specificity, and a truncated sequence can resemble a biologically generated HLA ligand. More generally, source-fragmented, proteolytically generated, or co-purified peptides may be misinterpreted as naturally presented ligands^30,31^. The explicit integration of pre-CID monitoring into our optimization therefore represents an important quality-control component. The apparent number of unique peptide sequences alone would have provided an incomplete and potentially misleading criterion for selecting the accumulation time.

Recent studies using DDA-PASEF, diaPASEF, and automated low-volume enrichment have demonstrated the potential of ion mobility-enabled workflows for limited samples.^16,20,32^ Nevertheless, absolute identification numbers from these studies are not directly comparable with those obtained here. Cell type, HLA genotype, HLA expression, biological state, starting material, immunoprecipitation efficiency of the used bead-antibody combination, injected fraction, chromatographic system, acquisition strategy, database search engine and specific settings, and criteria for defining an HLA ligand can all strongly influence the reported coverage. Tissue-dependent peptide yields and repertoire heterogeneity have also been observed even when samples are analyzed using the same platform.^31^ Consequently, the literature should be used to establish whether changing a configuration increases or decreases HLA-I identification within a controlled experiment, rather than to rank platforms solely according to peptide counts reported for different samples.

Computational rescoring is also a crucial improvement because acquisition sensitivity is useful only when spectra can be assigned confidently. Immunopeptidomics searches are intrinsically challenging because the absence of enzyme specificity substantially expands the sequence-search space. Fragmentation patterns of short, non-tryptic peptides also differ from those of the tryptic peptides on which many conventional scoring functions were developed. MS²Rescore^33^ addresses this problem by integrating search-engine scores with predicted fragment-ion intensities and chromatographic properties, while TIMS²Rescore^17^ additionally incorporates models adapted to timsTOF fragmentation and ion mobility. Importantly, recent benchmarking showed that rescoring gains are greater on newer-generation mass spectrometers, where improved sensitivity and acquisition speed provide more spectra that benefit from advanced rescoring^34^. This highlights the synergy between instrumental and computational advances. Its inclusion therefore complements the experimental optimization by improving discrimination between correct and incorrect peptide-spectrum matches rather than merely relaxing identification thresholds.

DDA-PASEF represents one component of a broader analytical landscape. Its precursor-level selectivity and interpretable MS/MS spectra support confident sequence assignment, while multiple search engines and data-driven rescoring provide additional options for assessing identification confidence.^32,35^ diaPASEF offers complementary gains in quantitative completeness and reproducibility across related samples but often relies on empirical or predicted spectral libraries. Although predicted libraries can include previously unobserved peptides, confidence depends on the accuracy and training coverage of HLA-binding and spectral-property models, particularly for novel or poorly represented immunopeptides^28,32^. Targeted approaches may be preferable for highly sensitive confirmation of predefined antigens^36^. These approaches should therefore be considered complementary and selected according to the biological question. Overall, the present work illustrates why no single acquisition configuration should be proposed as universally optimal and that sample-specific optimisations are often beneficial.

## Conclusion

In conclusion, this study establishes optimized DDA-PASEF-based immunopeptidomics workflows that improve HLA class I peptide identification, especially from limited sample material. By adapting a magnetic bead-based sample preparation protocol and carefully balancing ion accumulation time, number of PASEF scans, ion mobility range, collision energy ramps and customized polygon filtering, we enhanced sensitivity while minimizing pre-CID truncation and maintaining suitable throughput. The optimized sample preparation workflow enabled robust peptide recovery from substantially reduced starting material and improved compatibility with automation, supporting its potential application to clinical sample processing.

The optimized approach enabled broader detection of immunopeptides, including singly charged species that are often underrepresented in conventional methods. Together with TIMS²Rescore, this workflow provides a robust strategy for improving confidence and depth in immunopeptidome profiling. Overall, these methodological advances support the application of timsTOF-based immunopeptidomics to low-input and clinically relevant samples, while highlighting the benefits of fine optimization of both sample preparation and MS acquisition parameters, particularly on highly sensitive instruments such as the timsTOF Ultra.

## Methods

### HL60 cell culture

HL60 cell line was cultured in RPMI media supplemented with 10% fetal bovine serum and 1% Penicillin/streptomycin (Gibco) at 37°C with 5% CO2. Cells were grown up to 2e6 cells/mL and collected at 1500rpm. Cells were washed 3 times with ice-cold PBS and frozen at -80°C.

### B lymphocytes isolation

Blood samples were collected from CLL (Chronic Lymphocytic Leukemia) patients using 5-mL blood draws. Peripheral blood mononuclear cells (PBMCs) were isolated by density-gradient centrifugation using Ficoll™. B cells were subsequently enriched by immunomagnetic negative selection using the EasySep™ Human B Cell Enrichment Kit without CD43 (StemCell Technologies, #17963), which depletes non-B-cell populations using antibody-conjugated magnetic particles targeting unwanted cell populations. Enrichment purity was assessed by flow cytometry using antibodies against CD45, CD19, CD20, kappa, and lambda markers, yielding preparations containing >98% tumor B cells. Cells were washed 3 times with cold PBS and were subsequently stored at -80°C until further use.

### Immuno-affinity purification of HLA class I peptide ligands

#### Sepharose beads based sample preparation protocol

HLA class I ligands were enriched by immunoprecipitation as previously described^33^. A total of 600 x 10^6^ in-house cultured HL60 cells were lysed with 20mM Tris-HCl, 150mM NaCl, 0.25% sodium deoxycholate (m/v), 1mMEDTA pH8, 0.2mM iodoacetamide, 1mM PMSF, Roche Complete Protease Inhibitor Cocktail, 0.5%NP 40 (v/v), PBS, pH7,4. The lysate was centrifuged at 21,000×g for 30min at 4°C. HLA-peptide complexes were captured on CNBr-activated Sepharose 4B beads (Cytiva) linked to W6/32 antibody. Following binding, beads were washed several times with 3 buffers (150mMNaCl, 20mM Tris-HCl, pH 7.4; 400mM NaCl, 20mM Tris-HCl, pH 7.4; and 20 mM Tris-HCl, pH 8.0) and bound complexes were eluted in 0.1M acetic acid. Eluted HLA peptides and the subunits of the HLA complexes were desalted using a C18 Macro Spin column (Harvard Apparatus) according to the manufacturer’s protocol. Finally, HLA peptides were purified from the HLA-I complex after the elution with 25% ACN (v/v), 0.1% TFA (v/v). Samples were evaporated under vacuum and resuspended in water with 0.1% FA (v/v).

#### Magnetic beads based sample preparation protocol

HLA class I ligands were enriched by immunoprecipitation using an adapted protocol from *Liam Kam Sian et al.*, 2023. Only low-binding tubes were used throughout the experiments. The anti-pan HLA class I antibody W6/32 was purchased from Leinco Technologies (#H263). A total of 60 × 10⁶ in-house cultured HL60 cells were lysed in 200 µL of lysis buffer containing 20 mM Tris-HCl (pH 8.0), 150 mM NaCl, 0.5% Igepal, 0.25% sodium deoxycholate (w/v), 0.2 mM iodoacetamide, and 1/100 (v/v) protease and phosphatase inhibitors in PBS (pH 7.4). After one hour of rotation at 4°C at 25 rpm, the lysate was centrifuged at 20,000 × g for 10 minutes at 4°C, followed by a second centrifugation for 30 minutes.

HLA-peptide complexes were captured using magnetic beads (MagReSyn® Protein A Max beads, Resyn Biosciences). For each sample, 80 µL of beads were washed twice with 2 volumes (160 µL) of PBS and finally resuspended to the initial volume (80 µL). A total of 500 µg of W6/32 antibody was added to the beads and incubated with slow rotation (25 rpm) at 4°C for 30 minutes. After brief centrifugation, the bead-antibody complexes were washed twice with 1 mL of PBS and resuspended in 100 µL of PBS. These complexes were then added to the lysate and incubated overnight at 4°C with slow rotation (20 rpm). Following binding, beads were washed sequentially with three buffers: A (150 mM NaCl, 20 mM Tris-HCl, pH 8.0), B (400 mM NaCl, 20 mM Tris-HCl, pH 8.0), and C (20 mM Tris-HCl, pH 8.0), in the order A, B, A, C. Bound complexes were eluted twice using 200 µL of 0.1% TFA, with agitation at 1000 rpm for 20 minutes each time.

Eluted HLA peptides were desalted using a Sep-Pak tC18 cartridge (#WAT054960, Waters). The cartridge was conditioned with 2 mL of MeOH, 3 mL of ACN, and 3 mL of H₂O, 0.1% FA (v/v). Samples (400 µL) were loaded and washed with 2 mL of H₂O, 0.1% FA (v/v). Finally, peptides were eluted with 28% ACN (v/v) and 0.1% TFA (v/v). Samples were evaporated under vacuum, resuspended in 12 µL of H₂O, 0.1% FA (v/v). For DDM experiments, DDM was added at a final concentration of 0.015%. Samples were centrifuged at 17,000 × g at 4°C for 20 minutes. The supernatant was transferred to a new vial prior to nanoLC-MS/MS analysis.

### LC-MS/MS timsTOF Ultra2 settings

timsTOF Ultra (Bruker Daltonics, Bremen, Germany) was coupled to a nanoElute2 LC instrument (Bruker Daltonics). Peptides were separated by reversed-phase chromatography using a 25 cm Aurora nanoflow-packed emitter column (Ionoptics) featuring a 75µm inner diameter, 1.7µm particle size, and 120 Å pore size with a gradient of 2-35% B (2% ACN, 0.1% FA) over 60 or 100 minutes at a 0.2 µl/min flow rate. Buffer A consisted of 0.1% formic acid in water, while Buffer B contained 0.1% formic acid in acetonitrile.

The mass spectrometer was operated in positive ionization mode with Data-Dependent Acquisition Parallel Accumulation Serial Fragmentation (DDA-PASEF)^29^ using an HLA-tailored polygon. A full MS scan over a mass range of m/z 100-1700 and an ion mobility range from 0.60 to 1.7 1/K0 (V.s/cm^2^) were set, resulting in a cycle time of 1.45 s with 6 PASEF MS/MS ramps with charge states from 0 to 5+. Precursor ion selection width was kept at 2-Th@700 m/z and 3-Th @800 m/z. Ion accumulation and ramp time were set to 200 ms, and the collision energy was applied as a linear ramp from 20 eV at 1/K0 = 0.60 V.s/cm^2^ to 59 eV at 1/K0 = 1.6 V.s/cm^2^. MS2 scan range was set from 100 to 1700m/z; the target intensity value was set to 20.000 with an intensity threshold of 1500. Active Exclusion was enabled and set to 0.40 min.

Different LC (column size and gradient length) and MS parameters (accumulation times, collision energy, and number of PASEF scans) were evaluated. They are presented in the results section. The gradient used for LC method evaluation is presented in **Suppl. Table 2.**

### LC-MS/MS timsUltra AIP settings

The same LC and MS settings were used than previously described for timsTOF Ultra2. The Athena Ion Processor (AIP) was operated with a voltage ramp from 800 to 0 V, using a ramp slope of 30 V/µs for MS scans and 25 V/µs for MS/MS scans. The storage ramp was set to 0.7 1/K₀, with storage times of 12 µs for MS and 15 µs for MS/MS. The transfer time was set to 60 µs. All other parameters were not changed.

### Data processing pipeline and rescoring

Data analysis was performed on the Cumulus platform (https://github.com/LSMBO/cumulus-client, v2.2.5) using Sage software (v 0.15.0) and rescored with TIMS²Rescore^33^ (v 3.2.0). The protein human sequence database was downloaded from UniProtKB-SwissProt^37^ (organism ID 9606, 20,420 reviewed entries, downloaded on 25 July 2024). The common contaminants and decoy sequences were added before the search. The search was done with 10 ppm precursor and fragment tolerances and a maximum of two variable modifications. Oxidation of methionine, carbamidomethylation, acetylation and cysteinylation of cysteine were considered as variable modifications. No cleavage rule was used, and peptide lengths were set to 7−20. The data were rescored using TIMS²Rescore, with all feature generators enabled and the timsTOF2024 model. TIMS²Rescore leverages machine learning to predict retention times and fragment ion intensities via DeepLC^38^ and MS²PIP^39^, and integrates these with Mokapot^40^ for semi-supervised FDR estimation. The identification files were filtered to retain precursor charge at 1% FDR. MS1 quantification on the DDM dataset was performed using FlashLFQ GUI^41^ (v1.2.4) with the default settings and intensities were log2-transformed for visualization. Candidate pre-CID truncated peptides were identified within each LC–MS/MS run by comparing stripped peptide sequences after filtering target PSMs at a mokapot q-value ≤0.01. A peptide was classified as a truncated when its sequence exactly matched to another identified longer peptide after removal of one or two amino acids from either the N-or C-terminus.

## Acknowledgements

This work was supported by the Agence Nationale de la Recherche via the French Proteomic Infrastructure (ProFI UAR2048; ANR-10-INBS08-03 & ANR-24-INBS-0015) and by the Region Grand-Est (SC-Proteomics project) for equipment funds. J.B.R. and C.C. acknowledge CNRS and ANR for the ProteoVir project [ANR-24-RRII-0001]. J.B.R., C.K and C.C. acknowledge the Interdisciplinary Thematic Institute IMS, the drug discovery and development institute, as part of the ITI 2021-2028 program of the University of Strasbourg, CNRS and Inserm, supported by IdEx Unistra [ANR-10-IDEX-0002], and by SFRI-STRAT’US project [ANR-20-SFRI-0012]. A.D., R.G., C.C. and L.M. acknowledge funding from the Research Foundation Flanders (FWO) [1SE3724N, 12AK526N, G010023N, W005325N]. A.D. and L.M. acknowledge funding from the Horizon Europe Projects BAXERNA 2.0 [101080544] and COMBINE [101191739], and from the Ghent University Concerted Research Action [BOF21/GOA/033, BOF/PDO/2026/041]. C.C. and L.M. acknowledge funding from the CHIST-ERA project ODEEP-EU [ANR-23-CHRO-0005 and G0GDV23N]. Views and opinions expressed are however those of the author(s) only and do not necessarily reflect those of the European Union or the European Research Executive Agency (REA). Neither the European Union nor the granting authority can be held responsible for them.

## Supplementary Informations

**Suppl. Fig. 1.**
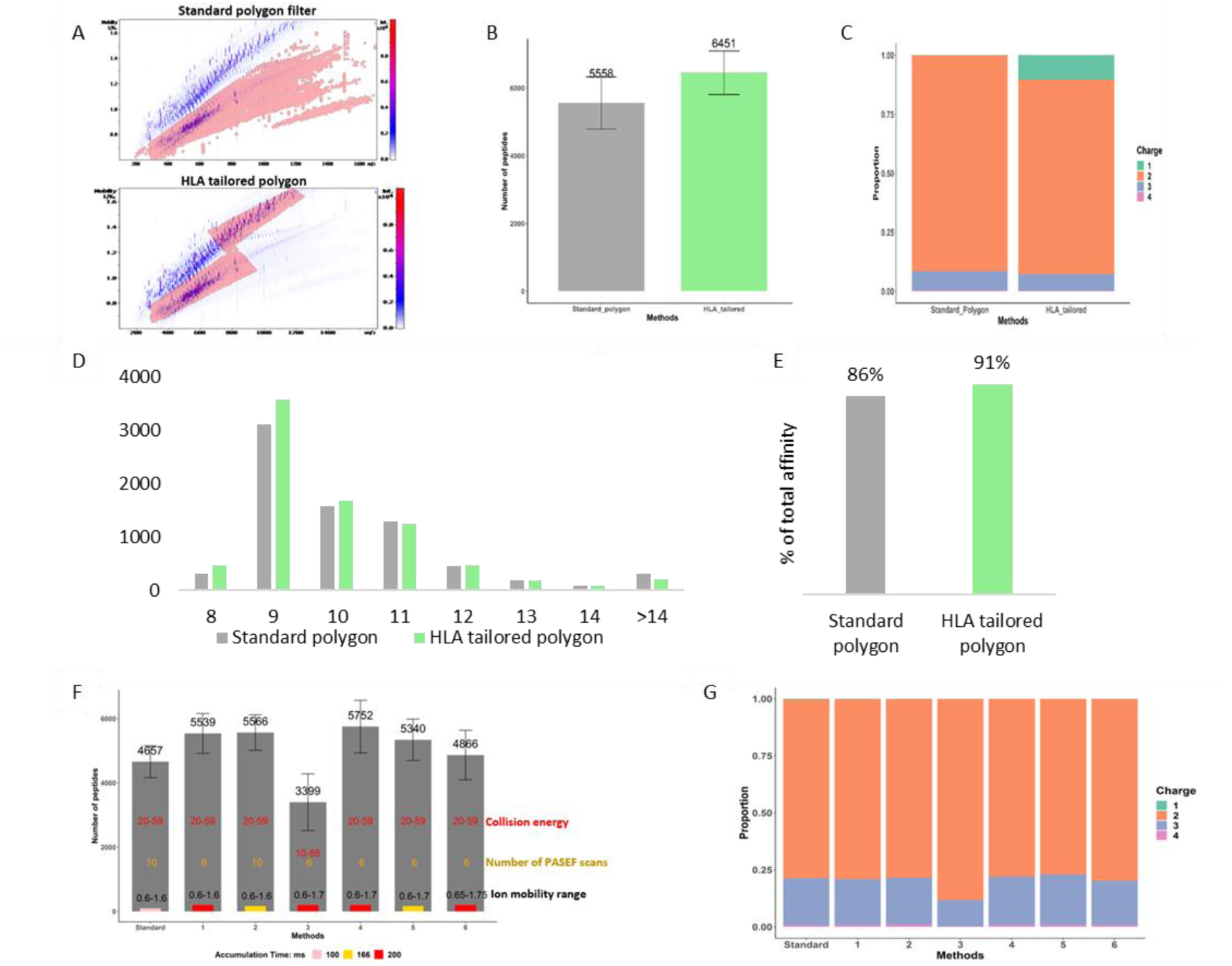
Method optimizations for immunopeptides on timsTOF Pro2. A) Design of the standard polygon filter (upper) and HLA-tailored polygon filter (lower) is shown in m/z vs IM plane. B) Number of peptides identified using the standard polygon filter and the HLA-tailored polygon filter. C) Peptide charge distributions using the standard polygon versus the HLA-tailored filter. D) Distribution of peptide length in standard and HLA-tailored polygon. E) Total binding affinity prediction for the 8-13 mers using netMHCpan 4.1 software for standard and HLA tailored polygon. F) Bar chart showing the number of peptides identified using different MS methods. Different parameters are represented with their respective colors. The accumulation time is represented by small inner bars: 100ms (pink), 166ms (orange), and 200ms (red). G) The proportion of differently charged peptide ions in the respective methods.

**Suppl. Table. 1.** Gradients used for LC methods evaluation

| **Gradient time** | **10 min** | **15 min** | **20 min** | **30 min** | **60 min** | **100 min** |
|---|---|---|---|---|---|---|
| **Total gradient time** | 16 min | 22 min | 30 min | 40 min | 80 min | 120 min |
| **2% ACN** | 0 min | 0 min | 0 min | 0 min | 0 min | 0 min |
| **23% ACN** | 8 min | 12 min | 16 min | 18 min | 36 min | 70 min |
| **35% ACN** | 10 min | 15 min | 20 min | 30 min | 60 min | 100 min |
| **90% ACN** | 14 min | 19 min | 26 min | 35 min | 70 min | 110 min |
| **90% ACN** | 16 min | 22 min | 30 min | 40 min | 80 min | 120 min |

**Suppl. Table. 2.** Total cycle time (in s) according to accumulation time and number of PASEF scans

| **Accumulation time (ms)** | **Number of PASEF scans** | **Total cycle time (s)** |
|---|---|---|
| 50 | 6 | 0.40 |
| 100 | 6 | 0.75 |
| 200 | 6 | 1.45 |
| 300 | 6 | 2.15 |
| 300 | 3 | 1.23 |

**Suppl. Fig. 3.**
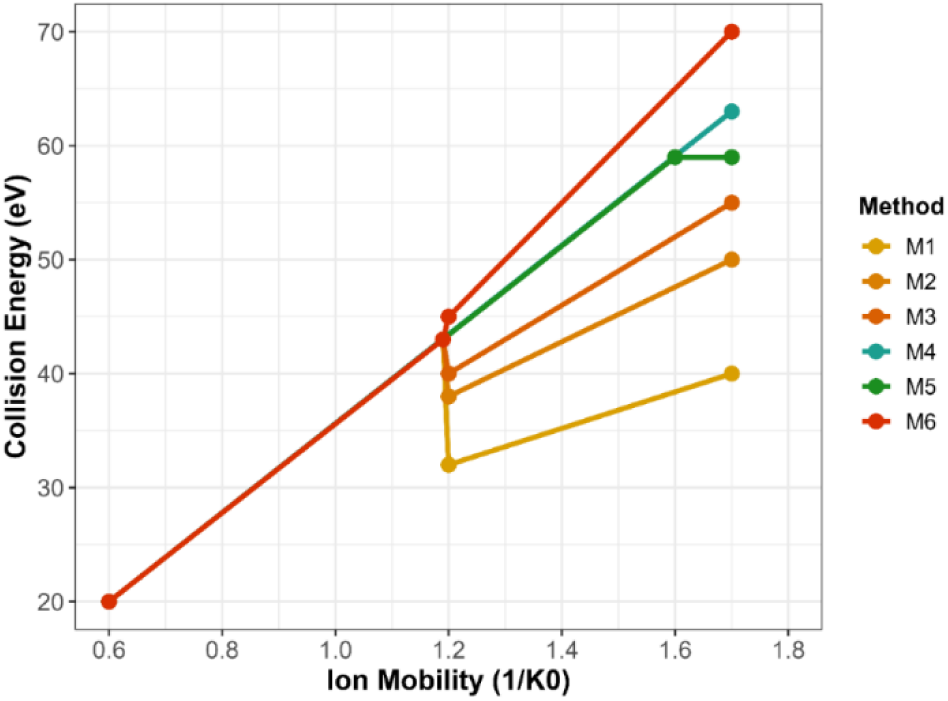
Different collision energy ramps (M1 to M6) used in the experiment to optimize the collision energies according to ion mobility range.

**Suppl. Fig. 4.**
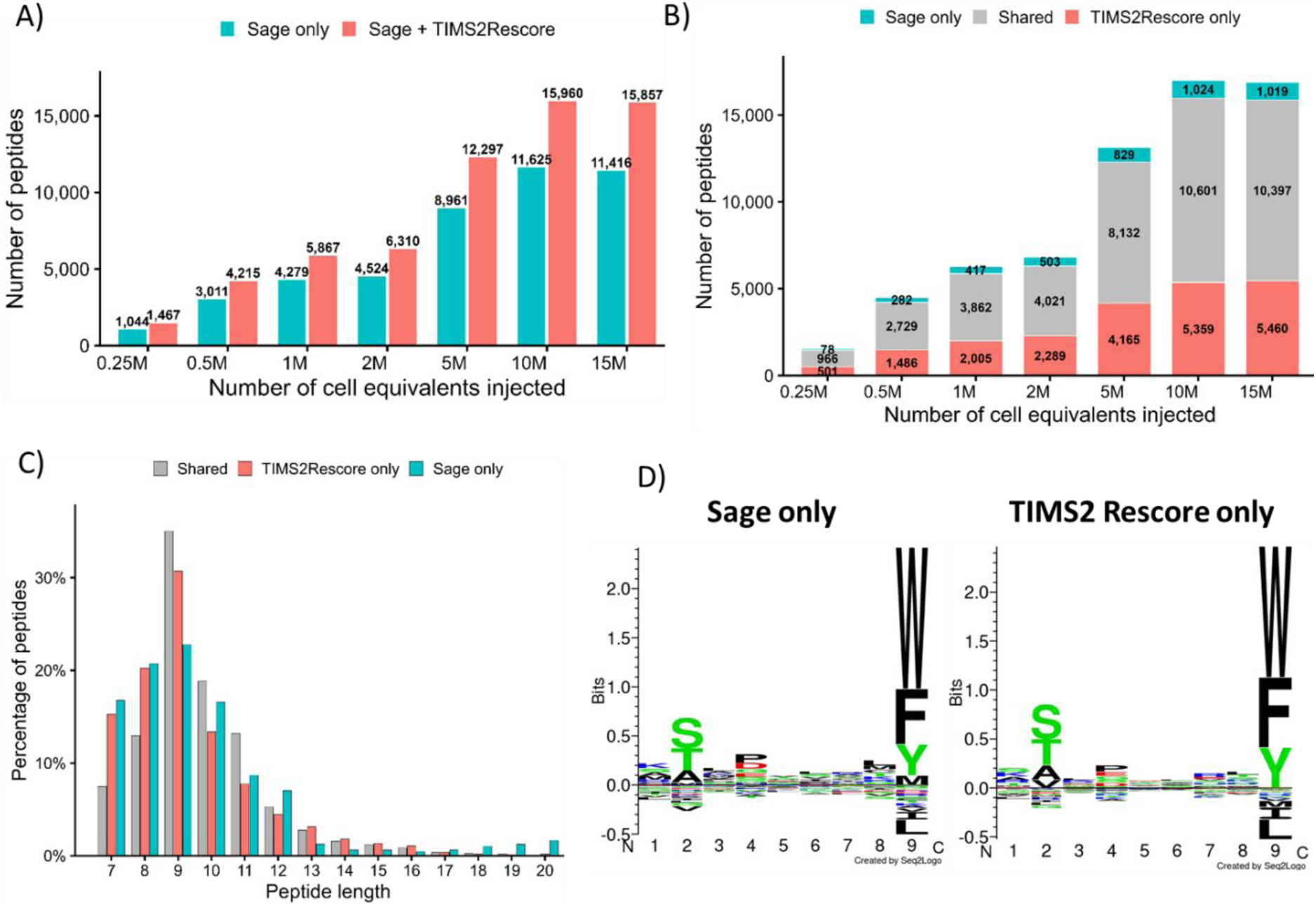
Peptide identification with the implementation of TIMS²Rescore. A) Number of peptides identified following injection of HLA peptides corresponding to 0.25–15 million HL60 cell equivalents using a 25 cm column and a 60-min LC gradient, with or without TIMS²Rescore. B) Number of peptides shared between Sage and TIMS²Rescore and uniquely identified by each approach. C) Peptide length distribution of shared and uniquely identified peptides. D) GibbsCluster motifs of peptides uniquely identified by Sage or by TIMS²Rescore.

